# VEGF-B/NRP1 Signaling Modulates Mitochondrial Homeostasis and Cardiac Function After Myocardial Infarction

**DOI:** 10.1101/2025.04.18.649613

**Authors:** Sai Manasa Varanasi, Ankit Sabharwal, Riya Kar, Carter Magnano, Ananya Dorairaj, Enfeng Wang, Shamit Dutta, Pritam Das, Stephen C Ekker, Ying Wang, Debabrata Mukhopadhyay, Ramcharan Singh Angom

## Abstract

**Background:** Myocardial infarction (MI) remains a leading cause of mortality worldwide. Recent studies suggest a cardioprotective role for vascular endothelial growth factor-B (VEGF-B) in MI. However, the molecular mechanisms of VEGF-B-mediated signaling via its co-receptor Neuropilin-1 (NRP1) in MI are poorly understood. In this study, we investigated the intricate signaling mechanisms involving VEGF-B and NRP1 in cardiomyocytes (CMs) using ischemic injury as a model of MI. And secondly, we further validated the protective role of VEFG-B in ischemic heart disease, shedding light on their roles not only in cardiac function but also in its therapeutic potential.

**Methods:** In this study, we utilized both *in vitro* and *in vivo* approaches to elucidate the role of VEGF-B and NRP1 signaling in MI and how it manifests a protective role in mitochondrial functions and cardiac regeneration following ischemic injury. We used two different cardiomyocyte cell lines, H9c2 (rat ventricular cardiomyocytes) and HL-1 (mouse ventricular cardiomyocytes), and induced hypoxia conditions, using either 1% oxygen or 200µM cobalt chloride (CoCl_2_) to mimic the myocardial infarction-induced ischemic injury in the heart. In addition, we developed a novel heat shock inducible zebrafish model of a cardiomyocyte-specific VEGF-B overexpression system to further examine the protective role of VEGF-B *in vivo*.

**Results:** Our findings indicate that both VEGF-B and NRP1 are predominantly expressed in heart tissue compared to other tissues, and their expression is altered in response to hypoxia/ischemic injury. Our results demonstrate that VEGF-B treatment prior to hypoxia enhances cardiomyocyte survival, while NRP1 knockdown abolishes this protective effect, highlighting a prominent role of NRP1 signaling in VEGF-B-mediated cardio protection. Furthermore, we found that VEGF-B promotes cardiomyocyte survival by improving mitochondrial function, as evidenced by reduced oxidative stress and ROS accumulation, decreased oxidative stress, preserved mitochondrial membrane potential, and increased ATP levels. Lastly, using our VEGF-B transgenic zebrafish model, we demonstrated that VEGF-B overexpression in zebrafish cardiomyocytes protects the heart from ischemic injury and enhances cardiac regeneration in an NRP1-dependent manner.

**Conclusion:** Our study has uncovered an important role of VEGF-B-NRP1 signaling axis in VEGF-B mediated cell survival and in beneficial mitochondrial functions in the CMs. Importantly, we also demonstrated that VEGF-B is protective against ischemic injury *in vivo* using a novel zebrafish model

**Novelty and Significance:** *What Is Known?:*

1. Myocardial infarction (MI) results in mitochondrial dysfunction, cardiomyocyte death, and adverse ventricular remodeling.
2. Vascular Endothelial Growth Factor B (VEGF-B) is traditionally associated with cardiac metabolism, survival, vascular biology, particularly in angiogenesis and lipid metabolism.
3. The neurophilin-1 (NRP1) receptor, a co-receptor for VEGF-B, is expressed in cardiomyocytes and implicated in mitochondrial signaling pathways.
4. Emerging evidence suggests that VEGF-B may influence mitochondrial integrity and function under stress conditions.

*What New Information Does This Article Contribute?:*

1. VEGF-B exerts a protective effect on cardiomyocyte mitocondria following ischemic injury by preserving mitochondrial function and reducing oxidative stress.
2. NRP1 receptor as a key mediator of VEGF-B modulates mitochondrial homeostasis in cardiomyocytes.
3. VEGF-B signaling can attenuate apoptosis and enhance mitochondrial bioenergetics post-MI, suggesting a novel cardioprotective mechanism.
4. Clinical Implication: These findings suggest that therapeutic strategies aimed at enhancing VEGF-B/NRP1 signaling may improve mitochondrial function and cardiac recovery after MI, offering a novel avenue for limiting heart failure progression in ischemic heart disease.

## Introduction

Cardiovascular diseases (CVDs) remain the leading cause of mortality worldwide, significantly impacting both physical and mental well-being^1,2^. The key risk factors contributing to CVDs include dyslipidemia, smoking, diabetes, and obesity^3^. Therapeutic interventions targeting these risk factors have been effective in reducing CVD-related morbidity and mortality^4^. Among CVDs, myocardial infarction (MI)—a severe manifestation of ischemic heart disease (IHD)—results from prolonged ischemia and hypoxia, leading to irreversible myocardial cell death^3^. Current clinical interventions for MI primarily focus on revascularization strategies, including percutaneous coronary intervention, thrombolysis, and coronary artery bypass grafting^5–7^. However, these approaches do not fully address post-MI cardiac remodeling, mitochondrial dysfunction, and cardiomyocyte death, necessitating the exploration of novel therapeutic strategies.

MI leads to structural cardiac remodeling, including scar tissue formation, cardiomyocyte hypertrophy, and microvascular dysfunction, all of which contribute to progressive heart failure^8^. The underlying pathophysiology of MI is driven by an imbalance in oxygen supply and demand, which triggers a cascade of cellular dysfunction, including oxidative stress, inflammation, and mitochondrial impairment ^3^. Recent studies have identified ferroptosis, an iron-dependent form of non-apoptotic cell death, as a major contributor to cardiomyocyte loss in MI^3,9^. Unlike apoptosis or necrosis, ferroptosis is characterized by mitochondrial shrinkage, depletion of glutathione, excessive lipid peroxidation, and accumulation of reactive oxygen species (ROS)^3,10,11^. These events exacerbate mitochondrial dysfunction and impair cardiac function, further worsening MI outcomes^12,13^.

Vascular endothelial growth factor B (VEGF-B), a key member of the VEGF family, plays a pivotal role in cardiomyocyte survival, vascular homeostasis, and mitochondrial function^14^. Unlike VEGF-A, which primarily promotes angiogenesis, VEGF-B is involved in mitochondrial protection, metabolic regulation, and endothelial cell function^15^ and survival^16^. Given its positive effects on angiogenesis and cardiac function, VEGF-B has been considered a potential therapeutic target for heart failure^17^. Although the potential therapeutic role of VEGF-B in heart failure remains promising, further research is needed to fully understand its mechanisms and to develop targeted interventions.

VEGF-B binds to VEGFR-1 and its co-receptor Neuropilin −1 (NRP1), leading to activation of downstream tyrosine kinase receptors such as p38 MAPK, ERK/MAPK, AKT, and PI3K^15,18^. NRP1 as a co-receptor for VEGF-B, forms a signaling complex with VEGFR-1 to activate pro-survival pathways that counteract ferroptosis and oxidative damage^15,18^. Beyond its role in VEGF-B signaling, NRP1 interacts with other molecular pathways that regulate cellular stress responses, endothelial barrier integrity, and remodeling of the extracellular matrix ^19,20^. Recent evidence suggests that NRP1 can enhance endothelial cell migration, promote vascular stability, and modulate immune cell infiltration in ischemic tissues, thereby contributing to improved cardiac repair post-MI ^21,22^. Furthermore, NRP1 has been linked to mitochondrial fission and fusion dynamics, suggesting its involvement in preserving mitochondrial integrity during hypoxic stress ^23^. These multifaceted roles of NRP1 underscore its potential as a therapeutic target in the treatment of cardiovascular disease. Additionally, in silico analysis of publicly available transcriptomic datasets has revealed consistent expression of VEGF-B and NRP1 in healthy human myocardial tissue. These findings further emphasize the significance of VEGF-B and NRP1 signaling as a potential therapeutic target for mitigating myocardial injury and promoting cardiac repair. In this paper, we further examined the molecular signaling pathways of the VEGF-B and NRP1 signaling axis in cell survival and mitochondrial function in CMs in response to hypoxia/ischemic injury and validated the VEGF-B-mediated protective responses against ischemic injury *in vivo* using a novel zebrafish model expressing VEGF-B in the heart.

## Materials and methods

### Ethical statement

All zebrafish experiments were conducted in accordance with the guidelines and regulations of the Institutional Animal Care and Use Committee (IACUC) at Mayo Clinic Jacksonville, Florida. Protocols were approved under animal use protocol number A00018914. Zebrafish (Danio rerio) were maintained under standard laboratory conditions in compliance with the Guide for the Care and Use of Laboratory Animals and the ARRIVE (Animal Research: Reporting of In Vivo Experiments) guidelines. Efforts were made to minimize animal suffering and to reduce the number of animals used.

### Data Availability

The data supporting this study’s findings are available from the corresponding author on reasonable request. Please see the Major Resources Table in the Supplemental Materials. The online Data Supplement presents an expanded version of the Methods section.

### Gene Expression Correlation Analysis Across Human Tissues

To investigate the correlation between NRP1 and VEGFB transcript expression across human tissues, we utilized publicly available RNA-sequencing data from the Genotype-Tissue Expression (GTEx) project (https://gtexportal.org/home/). Transcript per million (TPM) values for NRP1 and VEGFB were retrieved for all tissue types. Expression values were log-transformed prior to correlation analysis. Pearson correlation coefficients were calculated to assess the linear relationship between NRP1 and VEGFB expression levels across all tissue types, and scatter plots were generated to visualize the expression patterns.

### Cardiac Coexpression Analysis in Ischemic and Nonischemic Heart Tissue

To evaluate coexpression patterns of NRP1 and VEGFB in diseased versus healthy cardiac tissue, we used microarray data available in the iLINCS portal (Integrative LINCS genomics data portal; https://ilincs.org). Specifically, we analyzed data from 37 samples obtained using the Affymetrix Human Genome U133A Array (HG-U133A). The dataset included myocardial tissue samples from patients with ischemic cardiomyopathy (ICM) and nonischemic cardiomyopathy (NICM), two primary etiologies of dilated cardiomyopathy (DCM). Coexpression analysis was conducted separately for NICM and ICM groups to determine the relationship between NRP1 and VEGFB in each disease context. Expression normalization and differential correlation analysis were performed using the built-in statistical tools available within the iLINCS platform.

### Cell cultures

H9c2 embryonic rat heart-derived myoblast cells from ATCC and kindly donated by Dr. Nadine Norton at Mayo Clinic Jacksonville, were cultured in Dulbecco’s modified Eagle’s medium (DMEM), supplemented with 10% fetal bovine serum (FBS) under 95% air/5% CO_2_, and subculture when at 50–60% confluence. HL-1 cells from ATCC and a kind gift from Dr. Delisa Fairweather and Dr. Norton Nadine were grown in fibronectin–gelatine coated flasks containing Claycomb medium (“Sigma”) and supplemented with 10% FBS, 100 U/ml penicillin, 100 μg/ml streptomycin, 2 mM L-glutamine, and 0.1 mM norepinephrine. Cells were cultured at 37 °C and 5% CO_2_. Experiments were performed using cells between passages 8 and 12.

### Hypoxia Induction

H9c2 and HL-1 cells were subjected to hypoxia (1% O_2_) for different times (3h, 6h, 12h, 24h, 72h) in a controlled hypoxic plastic chamber (Stem Cell Technologies) and incubated at 37 °C respective times. Additionally, to mimic hypoxia, we also used 200µM cobalt chloride (CoCl_2_) and incubated cells at 37 °C for 24 to 72 hours as described before^24^.

### Data Retrieval and Preprocessing

The human heart tissue’s single-cell RNA sequencing (scRNA-seq) data were obtained from the Human Protein Atlas (HPA) database, which compiles data from published studies based on healthy human tissues^25^. This dataset comprises gene expression profiles for various tissues and cell types, with a primary focus on normal human heart tissue.

### Generating a Cardiac-Specific VEGF-B Expression Model in Zebrafish Using Heat Shock Induction

To generate a transgenic zebrafish model with cardiac-specific VEGF-B expression in AB strain under heat shock induction, we employed the strategy adapted from Hoeppner et al. (2012)^26^. A detailed protocol description is presented in the extended method as a supplementary file.

### Statistical analysis

All experimental data were analyzed using GraphPad Prism (Version 9) and are presented as mean ± standard error of the mean (SEM). Each experiment was performed in at least three independent biological replicates (n = 3) to ensure reproducibility. For comparisons between two groups, an unpaired two-tailed Student’s t-test was used. For multiple group comparisons, a one-way analysis of variance (ANOVA) followed by Tukey’s post hoc test was applied to determine statistical significance. For gene expression analysis via RT-qPCR, the ΔΔCt method was employed to calculate relative fold changes in mRNA levels, with target genes normalized to *β-actin* as the internal control. A p-value of less than 0.05 was considered statistically significant, with significance levels denoted as follows: *, p < 0.05; **, p < 0.01; **, p < 0.001; and ****, p < 0.0001. Data were visually represented using bar graphs or scatter plots, with error bars indicating the Mean ± standard deviation (SD).

## Results

### Correlation Analysis of NRP1 and VEGF-B Expression in Human Tissues

To investigate the relationship between NRP1 and VEGF-B expression across human tissues, a Pearson correlation analysis was performed using RNA-seq data from the GTEx database. Expression levels (TPM, transcripts per million) of NRP1 and VEGF-B were extracted for the normal human tissue types. The analysis revealed a significant positive correlation between NRP1 and VEGF-B expressions across the dataset as shown by (Pearson correlation coefficient r = 0.64 (p < 0.0001) as shown in the scatter plot (**Supplementary Fig. S1A**). These findings support our hypothesis that NRP1 and VEGF-B may be co-regulated or functionally linked in various tissues, including the heart. To assess the disease-contextual relationship between NRP1 and VEGF-B expression in the human heart, we performed coexpression analysis using transcriptomic data from the iLINCS portal^27^ https://www.ilincs.org/ilincs/. Coexpression analysis was conducted independently for the ischemic cardiomyopathy (ICM) and nonischemic cardiomyopathy (NICM) groups following data normalization using the platform’s integrated preprocessing pipeline. In both groups, NRP1 and VEGF-B transcripts exhibited a positive correlation **(Supplementary Fig. S1B)**. However, the strength of this correlation was notably higher in the ICM cohort, suggesting a potentially enhanced functional relationship under ischemic conditions. Differential correlation analysis confirmed a statistically significant increase in NRP1–VEGF-B coexpression in ICM compared to NICM myocardial tissue. These findings indicate that ischemic remodeling may involve coordinated upregulation or coregulation of NRP1 and VEGFB, supporting a context-specific role for this signaling axis in myocardial stress adaptation.

### Effects of VEGF-B in Promoting Cardiomyocyte Survival by Enhancing Proliferation

To determine the effect of hypoxia (1%O_2_ or CoCl_2_) on H9c2 and HL-1 cell proliferation, viability, and mitochondrial function, we treated these cells with hypoxia for 72 hrs to mimic ischemic injury in the MI. Normoxia-treated cells were used as controls. Ki67 immunostaining revealed that 72 hrs of hypoxia (1%O2) significantly reduced Ki67 expression in H9c2 cells (∼50% reduction compared to normoxic controls) **(Fig. 1A, 1B).** Similar results were observed for the HL-1 cells treated with 200 μM CoCl_2_ for 72hrs displayed reduced Ki67 positive cells (∼35%) as shown in (**Supplementary Fig. S2A, S2B**). RT-qPCR analysis confirmed a decrease in *Ki67* and *cyclin d1* mRNA levels (∼50% for *ki67* and 30-40 % for *cyclind1* in H9c2 cells treated with hypoxia for 24 hrs (**Fig 1C, 1D)** and in HL-1 cells treated with CoCl_2_ for 24 hrs (**Supplementary Fig. S2C, S2D)** compared to normoxia control. We then examined the effect of VEGF-B treatment on hypoxia-induced cell death in both the H9c2 **(Fig 1A-D)** and HL-1 cells **(Supplementary Fig S2 A-D)** by treating these cells with recombinant VEGF-B (20ng/ml), and hypoxia for 72 hr. VEGF-B treatment increased Ki67 staining in the H9c2 cells treated with hypoxia compared to those treated with hypoxia alone **(Fig. 1A, 1B).** We also observed an increased mRNA expression of *Ki67* and *cyclinD1 in the cells treated with VEGF-B and hypoxia compared to the hypoxia alone treatment* **(Fig. 1C,1D).** These results suggest that VEGF B ameliorates the H9c2 cell proliferation under hypoxia. Henceforth, we used 200 µM CoCl2 for all downstream experiments to induce hypoxia to mimic ischemic injury (MI) as reported earlier^28^. We then performed MTT assays to assess cell survival in both H9c2 and HL-1 cells treated with hypoxia (CoCl_2_), VEGF-B, and a combination of VEGF-B and hypoxia (CoCl_2_). CoCl_2_ treatment for 72 hrs resulted in a significant decrease in cell viability (∼40% reduction compared to normoxia) as indicated by a decrease in absorbance, reflecting hypoxia-induced cellular stress **(Fig. 1E, 1F).** Combination of VEGF-B and hypoxia-treatment showed partial recovery in cell viability compared to hypoxia-only treatment, indicating that VEGF-B ameliorates hypoxia-induced cell death in both H9c2 **(Fig. 1E),** and HL-1 cells **(Fig. 1F).**

**Figure 1.**
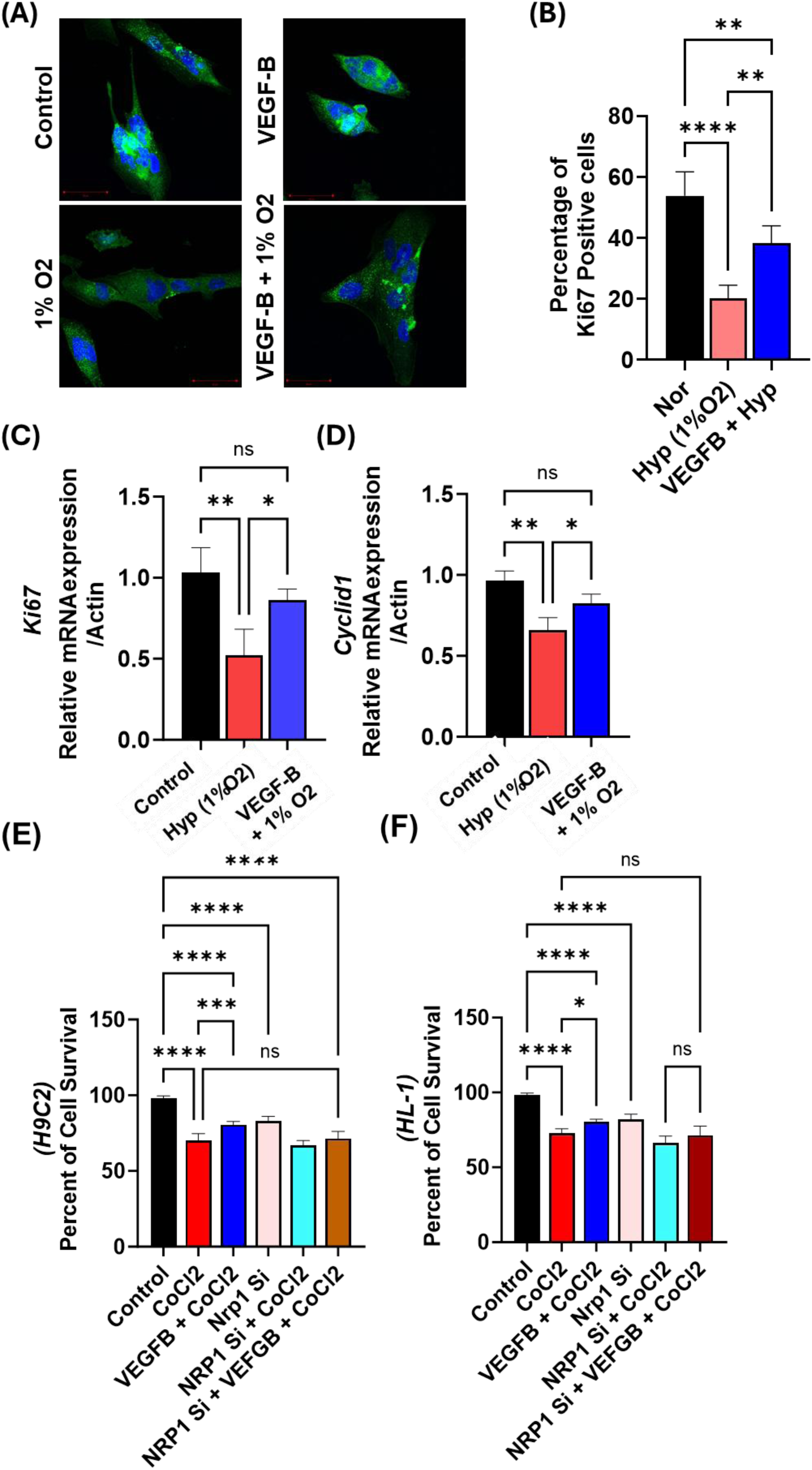
VEGFB Promotes Cardiomyocyte Survival by Enhancing Proliferation. **(A**) Immunofluorescence images showing Ki67 staining in cardiomyocytes (CMs) treated under normoxic (21% O₂) and hypoxic (1% O₂) conditions. **(B)** Quantification of Ki67-positive cells.. **(C)** mRNA expression analysis of Ki67 in H9C2 cardiomyocytes. **(D)** mRNA expression analysis of Cyclin D1 in H9C2 cardiomyocytes. **(E)** MTS assay showing VEGFB-mediated prevention of H9C2 CM death. **(F)** MTS assay showing VEGFB-mediated prevention of HL-1 CM death. Bars represent mean ± standard deviation (SD). Each experiment was repeated at least three times. Statistical significance: * p < 0.05. ** p < 0.01. ***, p < 0.001 and ****, p < 0.0001.

Further, to investigate the role of NRP1 in VEGF-B-mediated protection, we performed NRP1 knockdown in H9c2 and HL-1 cells by using NRP1 siRNA for 24 hours before treating them with hypoxia (CoCl_2_) alone and both CoCl_2_ and VEGF-B (20 ng/ml) together. NRP1 knockdown by itself reduced the viability (10-15%) **(Fig. 1E, 1F)**. NRP1 knockdown significantly diminished the protective effects of VEGF-B under hypoxia conditions, with cell viability in NRP1 knockdown cells treated with VEGF-B and hypoxia still comparable to that in hypoxia-only conditions in both H9c2 **(Fig. 1E)** and HL-1 cells **(Fig.1F)**. These results suggest that VEGF-B-mediated protection in hypoxic cardiomyocytes is dependent on NRP1.

### VEGF-B Activates Pro-Survival Signaling Pathways under Hypoxic Stress via NRP1

We next examined the effect of VEGF-B treatment on NRP1-mediated signaling in both H9c2 cells and HL-1 cells treated with hypoxia (200 μM CoCl2), VEGF-B alone, and a combination of VEGF-B and hypoxia. We first assessed protein levels of AKT, phosphorylated AKT (pAKT), ERK, phosphorylated ERK (pERK), cleaved caspase-3, caspase 3, Beclin-1, GPX4, which are involved in cell survival^29,30^, apoptosis^31^, autophagy pathways^32^, and other cell death pathways^33^. In control cells, basal expression levels of AKT, pAKT, ERK, pERK, caspase 3, cleaved caspase 3 **(Fig. 2A-D), and Bcl-2, Bax (Fig. S3A)** were readily detected. Upon hypoxia treatment, there was a significant decrease in the expression of pAKT and pERK protein levels as shown by western blot of whole cell lysates of H9c2 cells, and quantification of fold change for pAKT/AKT and pERK/ERK levels (**Fig. 2A-C)**. And this effect was more pronounced in HL-1 cells, which also show significantly reduced expression levels of pAKT and pERK protein levels after hypoxia **(Supplementary Fig. S3B, S3C)**, again indicating diminished activation of these key survival pathways following hypoxia treatment in H9c2 cells **(Fig. 2A and Supplementary Fig. S3A).** We observed significant upregulation of cleaved caspase 3 in the hypoxia treated cells **(Fig. 2A, 2D)** suggesting hypoxia-induced activation of apoptotic cell death pathways. Similar results were observed in HL-1 cells, for instance ratio of Bcl-2/Bax was also shifted toward pro-apoptotic signaling, with increased Bax expression and reduced Bcl-2 after hypoxia treatment **(Fig. S3)** and increased cleaved caspase 3 level further suggesting activation of apoptosis pathways in the CMs following hypoxia treatment **(Supplementary Fig S3B, S3C and S4).**

**Figure 2.**
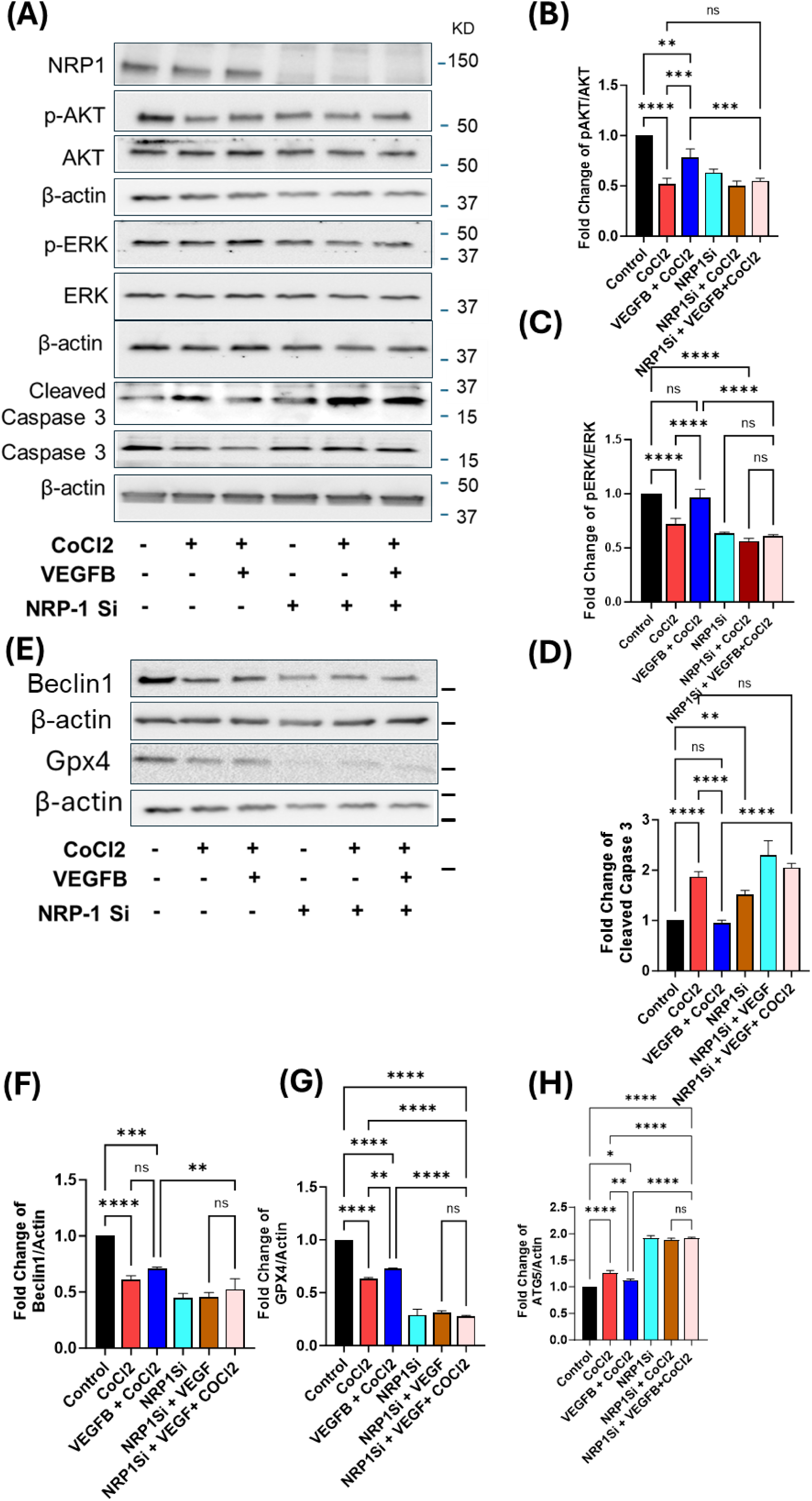
NRP-1 Knockout Inhibits the Protective Effects of VEGFB on Cell Survival and Protein Expression in H9C2 Cardiomyocytes. **(A)** Western blot analysis showing protein expression of cell survival and apoptotic pathway proteins in lysates from H9C2 cardiomyocytes (CMs) treated with various experimental groups.. **(B-D)** Quantification of the protein levels shown in (A). **(E)** Western blot analysis showing the expression of apoptosis markers and autophagy proteins in H9C2 CM treated with CoCl2 and NRP1siRNA with and without VEGFB. **(F-H)** Quantification of the Western blot analysis shown in (E), displaying fold changes in protein expression of the indicated markers. Each experiment was repeated at least three times. The bars represent the mean ± SD. Statistical significance is indicated as *p < 0.05, **p < 0.01, ***, p < 0.001, and ****p < 0.00001.

We then examined the effect of VEGF-B treatment on these signaling pathways in the CM’s. Treatment with VEGF-B under hypoxia conditions in H9c2 cells increased pAKT and pERK suggesting activation of cell survival signaling pathways in the VEGF-B-treated cells **(Fig. 2A-C)**. Similarly, treatment with VEGF-B in HL-1 cells showed both increased AKT and ERK and pAKT and pERK levels, confirming activation of cell survival pathways in the VEGF-B treated cells **(Supplementary Fig. S3**). Notably, Cleaved caspase-3 levels were also reduced in the H9c2 cells and HL-1 treated after VEGF-B, suggesting that VEGF-B protected the CMs from hypoxia-induced cell death **(Fig. 2A, 2D and Fig. S4)**. Furthermore, Bcl-2 protein levels were upregulated and Bax expression was reduced in the VEGF-B-treated HL-1 cells under hypoxia conditions, indicating a shift towards cell survival and anti-apoptotic signaling in HL-1 cells **(Supplementary Fig. S3)**. We first confirmed the NRP1 knockdown in the H9c2 cells as shown by reduced protein expression as measured by western blot (**Fig. 2A**). In NRP1 knockdown cells under hypoxic conditions, the levels of pAKT and pERK were reduced significantly considerably compared to VEGF-B-treated cells, indicating that NRP1 is crucial for VEGF-B-induced activation of these survival pathways **(Fig. 2A-2C)** and furthermore, in NRP1 knockdown cells treated with VEGF-B and hypoxia, the pAKT and pERK levels remained unchanged when compared to hypoxia-only treated cells, confirming that the protective effects of VEGF-B were lost when NRP1 was knocked down. The cleaved caspase-3 expression was increased in all groups of cells treated with NRP1 siRNA, indicating that NRP1 knockdown by itself activates pro-apoptotic pathways, and VEGF-B treatment in NRP1 knock out cells did not reduce/affect cleaved caspase activation after hypoxia treatment, suggesting that NRP1 knockdown abrogates the anti-apoptotic effects of VEGF-B in H9c2 cells **(Figs. 2A-2D)**.

Furthermore, to investigate the role of autophagy in hypoxia-induced cell death previously reported in other cell types^34^ we analyzed the protein levels of autophagy-related markers, including Beclin 1 and GPX4, in mitochondrial protein lysates isolated from H9c2 cardiomyocytes. Since mitochondria are both targets and regulators of autophagy, we focused on mitochondrial fractions to assess how mitochondrial autophagy, or mitophagy, might be affected under hypoxia. In parallel, we investigated the potential involvement of ferroptosis, a regulated form of cell death characterized by iron-dependent lipid peroxidation. GPX4 (glutathione peroxidase 4) is a mitochondrial antioxidant enzyme that prevents lipid peroxidation and serves as a central suppressor of ferroptosis. We isolated mitochondria from H9c2 cells and measured GPX4 protein levels to examine this. We observed a significant reduction in mitochondrial GPX4 expression in response to hypoxia **(Fig. 2E and G)**, indicating activation of ferroptosis pathways in cardiomyocytes following hypoxic stress. Collectively, the downregulation of Beclin 1 and GPX4 suggests that both autophagy dysregulation and ferroptosis may contribute to hypoxia-induced cardiomyocyte injury. As shown in figure **Fig. 2E, 2F**, we observed decreased expression of both Beclin1 and GPX4 in isolated mitochondrial under hypoxia conditions. This result indicates a possible role of ferroptosis mechanism in the cell death. In cells treated with VEGF-B along with hypoxia, we observed slightly reduced expression of GPX4 compared to the control group. However, VEGF-B treatment did not alter Beclin-1 protein levels **(Figs. 2E, 2F)**, suggesting that the autophagy pathways are differentially regulated. We next analyzed the effects of NRP1 knockdown on the VEGF-B-mediated signaling pathways in the CMs. Interestingly, we observed that in NRP1-depleted cells, the protein levels of both Beclin 1 and GPX4 were decreased **(Figs. 2E–2G),** indicating a compromised cellular capacity to manage stress through autophagy and ferroptosis suppression. The concurrent downregulation of these proteins suggests that NRP1 may be pivotal in regulating pathways that protect cardiomyocytes under stress conditions. Surprisingly, these levels were not altered in the control cells after hypoxia alone, VEGF-B alone or both hypoxia and VEGF-B treatment, suggesting that NRP1 knockdown interferes with VEGF-B’s ability to promote autophagy effectively. These results collectively demonstrate that VEGF-B treatment activates survival signaling through AKT and ERK, prevents apoptosis by modulating the Bcl-2/Bax ratio and inhibiting caspase-3 cleavage, and promotes autophagy through Beclin-1 and GPX4 expression in the H9c2 cells. Notably, the protective effects of VEGF-B were significantly diminished when NRP1 is knocked down, highlighting the critical role of NRP1 in mediating VEGF-B’s protective effects against hypoxia-induced stress in H9c2 cells and HL-1cells.

We then further validated the protective effect of VEGF-B and NRP1 signaling on hypoxia-induced apoptosis in H9c2 and HL-1 cardiomyocytes using the Apoptosis/Necrosis Detection Kit (Abcam, Cat# ab176749). Following treatment, cells were stained with Hoechst 33342 (blue, nuclear stain), Apopxin Green Indicator (early apoptosis marker), and 7-AAD (red, necrosis/late apoptosis marker), and analyzed by fluorescence microscopy **(Fig. 3A)**. We observed distinct differences among the experimental groups in H9c2 **(Fig. 3A)** and HL-1 **(Supplementary Fig. S5A, S5B)**. The control groups with and without VEGF-B showed minimal Apopxin-positive cells, strong Hoechst staining was observed with minimal Apopxin Green or 7-AAD fluorescence, indicating a predominantly viable cell population, and a low baseline level of apoptosis or necrosis **(Fig. 3A)**. In contrast, the hypoxia (CoCl_2_)-treated group showed a significant increase in Apopxin-positive and 7-AAD-positive cells **(Fig. 3A-C),** indicating elevated apoptosis or necrosis under hypoxic stress. This increase was further exacerbated in the NRP1 knockdown H9c2 group subjected to hypoxia **(Fig. 3A-C)**, where Apopxin staining showed markedly higher apoptotic cell counts. Interestingly, in the NRP1 knockdown cells treated with hypoxia and VEGF-B, Apopxin positivity remained elevated and comparable to NRP1 knockdown cells treated with hypoxia **(Fig. 3A, 3C),** suggesting a loss of protective signaling mediated by NRP1. This result highlights that VEGF-B’s protective effect against apoptosis is compromised in the absence of NRP1, reinforcing the essential role of NRP1 in VEGF-B-mediated cell survival.

**Figure 3.**
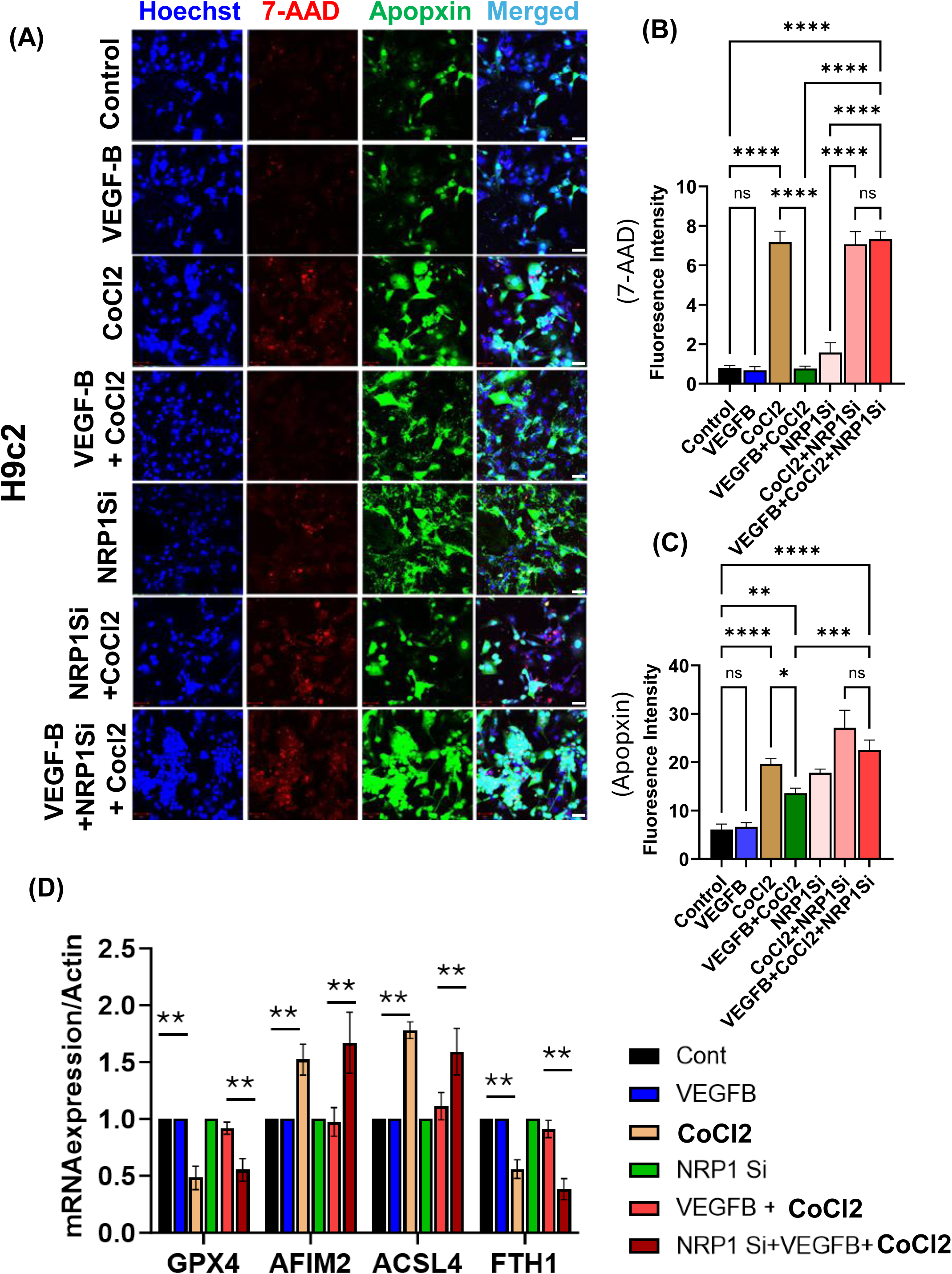
VEGF-B Prevents Cardiomyocyte Apoptosis through NRP1. **(A)** Representative immunofluorescence images of H9C2 cardiomyocytes (CMs) stained with Annexin V in Untreated/Control H9C2 cells, H9C2 cells treated with VEGF-B, H9C2 cells exposed to hypoxia by treating with 200 µM CoCl₂ for 72 hours, H9C2 cells treated with VEGF-B plus CoCl₂, H9C2 cells with NRP1 siRNA-mediated knockdown, H9C2 CMs with NRP1 knockdown plus CoCl₂ and H9C2 CMs with NRP1 knockdown plus CoCl₂ and VEGF-B treatment. **(B)** Quantifications of PI and **(C)** Quantification of Annexin V. **(D)** RT PCR analysis of ferroptosis marker genes. Bars represent mean ± standard deviation (SD) from three experiments. Statistical significance is indicated as *, p < 0.05, **p < 0.01, ***p < 0.001, and ****p < 0.0001. Experiments in each group were repeated at least three times. The images were captured using an LSM 800 confocal microscope with 488 nm and 594 nm lasers. Scale bar = 100 µm.

### Ferroptosis Pathway Activation in Hypoxia-Induced Cardiomyocyte Death and Its Modulation by VEGF-B and NRP1

Recent studies have identified ferroptosis, an iron-dependent form of non-apoptotic cell death, as a major contributor to cardiomyocyte loss in MI^3,9^. To investigate the role of ferroptosis in hypoxia-induced cardiomyocyte death and proliferation, we analyzed the mRNA expression of key ferroptosis-related genes in H9c2 cardiomyocytes exposed to hypoxia (CoCl2), VEGF-B, NRP1siRNA, and NRP1 siRNA plus Hypoxia plus VEGF-B. QRT-PCR analysis revealed that hypoxia significantly upregulated the expression of ferroptosis markers, including Acyl-CoA Synthetase Long-Chain Family Member 4 (ACSL4) which promotes ferroptosis by producing PUFA-containing phospholipids, making membranes more susceptible to lipid peroxidation^36^, Apoptosis-Inducing Factor Mitochondria-Associated 2 (AIFM2) which inhibits ferroptosis by regenerating CoQ10 and scavenging lipid radicals, independently of GPX4^37^, downregulation of Ferritin Heavy Chain 1 (FTH1) which protects against ferroptosis by storing free iron and preventing iron-mediated lipid ROS generation^38^ and Glutathione Peroxidase 4(GPX4) which regulate detoxification of lipid hydroperoxides and prevents ferroptosis, compared to control cells39,40 (p < 0.01) **(Fig. 3D)**, suggesting enhanced lipid peroxidation and iron-dependent cell death. Additionally, TFRC (transferrin receptor), a key regulator of iron uptake, was significantly increased in hypoxic cardiomyocytes, further supporting the activation of ferroptosis. Conversely, the reduced expression of GPX4 and FTH1, crucial inhibitors of ferroptosis, was markedly downregulated in hypoxic conditions (p < 0.01) **(Fig. 3D)**, indicating a loss of antioxidant defense and iron storage regulation. Treatment with VEGF-B under hypoxia partially suppressed ferroptosis, as evidenced by reduced ACSL4 and AIFM2 expression and increased GPX4 and FTH1 levels compared to hypoxia alone (p < 0.05) **(Fig. 3D)**. However, NRP1 knockdown in VEGF-B-treated hypoxic cells reversed these protective effects, leading to a sustained increase in ACSL4, AIFM2, expression, along with a significant decrease in GPX4 and FTH1 mRNA levels (p < 0.01 vs. VEGF-B plus hypoxia group). These findings suggest that hypoxia promotes cardiomyocyte ferroptosis by upregulating lipid peroxidation and iron regulatory pathways, while VEGF-B exerts a protective effect by downregulating ferroptosis-associated genes. The loss of NRP1 negates VEGF-B-mediated protection, highlighting the NRP1-dependent regulation of ferroptosis in hypoxic cardiomyocytes.

### Mitochondrial Function, ROS Generation, and Cellular Energy Homeostasis in Response to Hypoxia and VEGF-B Treatment

To determine mitochondrial function and reactive oxygen species (ROS) production in H9c2 and HL-1 cells following hypoxia and VEGF-B treatment, we utilized JC-1 staining, to measure the effects of VEGF-B and hypoxia treatment in studying the mitochondrial membrane potential (Red fluorescence -aggregate and green fluorescence-monomer). In control cells, aggregate JC-1 staining showed strong red fluorescence, indicating intact mitochondrial membrane potential (ΔΨm), with minimal green fluorescence, suggesting low levels of monomeric JC-1 **(Fig. 4A).** The NRP1 knockdown was confirmed by RT PCR analysis **(Fig. 4B).** VEGF-B treatment alone didn’t change the red fluorescence, indicating no change in the membrane potential **(Fig. 4A).** In contrast, hypoxia-treated cells exhibited a significant decrease in aggregate JC-1 levels (red fluorescence), indicative of mitochondrial depolarization and disrupted ΔΨm **(Fig. 4A, 4C)**. VEGF-B treatment along with hypoxia demonstrated a partial restoration of aggregate JC-1 levels (red fluorescence) when compared to hypoxia alone (**Fig. 4A, 4C),** suggesting that VEGF-B may offer a protective effect on mitochondrial membrane potential under hypoxic conditions.

**Figure 4.**
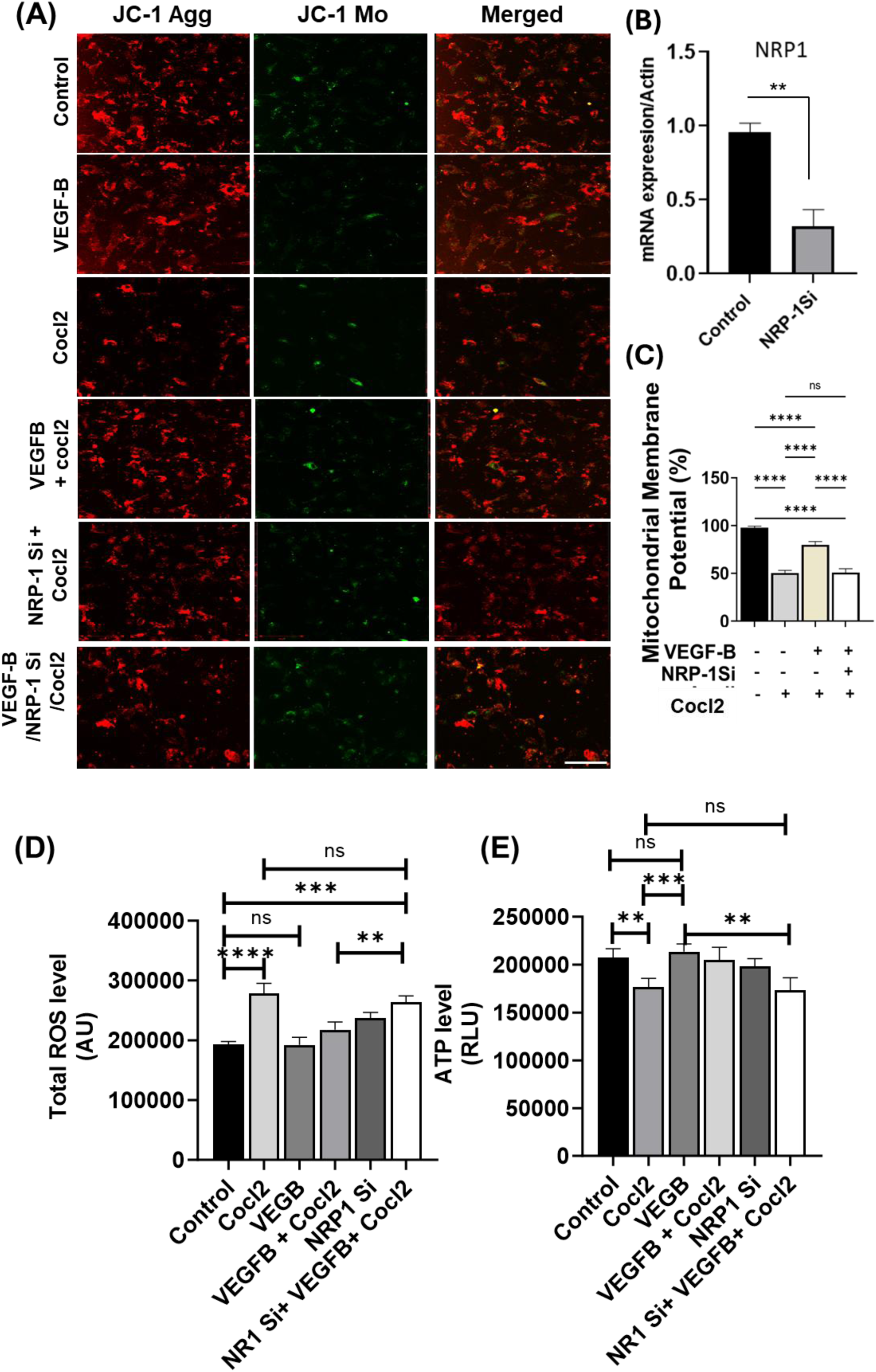
VEGF-B mitigates the mitochondrial function in ischemic CM through NRP1. **(A)** Representative confocal image showing H9c2 CMs stained with JC-1 dye after different exposures. **(B)** mRNA expression showing NRP1 knockdown in the CM. **(C)** Quantification of the mitochondrial membrane potential based on JC-1 staining. **(D)** Total ROS and **(E)** ATP levels were measured in the CMs after the treatments. Each experiment was repeated two times. The bars represent mean ± SD. *, p<0.05, **,p<0.01, ***, p<0.001, ****, p<0.0001.

We then examine the effects of VEGF-B and NRP1 on mitochondrial membrane potential in H9c2 cells after NRP1 ablation using siRNA-mediated NRP1 knockdown. NRP1 knockdown itself reduces the red fluorescence, suggesting altered membrane potential **(Fig. 4A).** SiRNA mediated NRP1 depletion prevented the protective effect of VEGF-B on mitochondrial membrane potential in the H9c2 cells, as the JC-1 intensity doesn’t change upon treatment with VEGF-B and hypoxia **(Fig. 4A, 4C**), confirming the critical role of NRP1 in the VEGF-B-mediated beneficial effects in CMs in response to hypoxia. To further examine the effects of hypoxia and VEGF-B on mitochondrial functions, we next measured mitochondrial ROS production in the CMs in response to hypoxic conditions and VEGF-B treatment using fluorescence dye DCFH-DA (10 µM) from Abcam #ab113851). Control H9c2 cells exhibited minimal mitochondrial ROS production, as indicated by basal DCF (Dichlorofluorescein) fluorescence level **(Fig 4D).** However, cells treated with hypoxia showed a significant increase in DCF fluorescence compared to the control **(Fig. 4D)**, indicating elevated production of ROS. Importantly, in H9c2 cells treated with VEGF-B under hypoxia reduced DCF fluorescence levels, suggesting an attenuation of oxidative stress **(Fig 4D**). VEGF-B treatment, however, was unable to reduce the mitochondrial ROS levels when NRP1 was abrogated using NRP1 siRNA when compared to hypoxia alone, indicating that VEGF-B may mitigate mitochondrial oxidative stress via NRP1. To further confirm these results, we measured Intracellular reactive oxygen species (ROS) levels in H9c2 cells were assessed using the CellROX Assay (Green reagent) for detection of oxidative stress. Following hypoxia treatment, a marked increase in CellROX fluorescence intensity was observed compared to control cells **(Supplementary Fig S6A, S6B)**, indicating elevated ROS production after hypoxia treatment. Quantitative analysis showed a significant increase in mean fluorescence intensity in treated groups (p < 0.01), consistent with enhanced oxidative stress (**Fig. S6**). These results demonstrate that CoCl_2_ induces oxidative stress in cardiomyocytes. However, VEGF-B treatment was unable to reduce the mitochondrial ROS levels when NRP1 was abrogated using NRP1 siRNA compared to hypoxia alone, indicating that VEGF-B may mitigate mitochondrial oxidative stress via NRP1 as observed above.

And lastly, to investigate the effects of VEGF-B treatment on cellular energy metabolism under stress conditions, ATP levels were measured using Realtime-Glo Extracellular ATP Assay (Promega#GA5010) in H9c2 cells treated with hypoxia and VEGF-B. Control cells maintained stable ATP levels, reflecting a normal cellular energy status **(Fig 4E).** However, hypoxia treatment significantly decreased ATP levels in the H9c2 cells **(Fig 4E),** indicating impaired mitochondrial function and cellular energy depletion under low-oxygen conditions. In contrast, VEGF-B treatment partially restored ATP levels in H9c2 cells treated with hypoxia compared to the untreated hypoxia-only group **(Fig. 4E)**, again suggesting that VEGF-B treatment helps preserve ATP production under hypoxic stress. These results highlight VEGF-B’s protective role in both H9c2 cells in response to hypoxic stress (need figure legend to write result for siNRP1), as it preserves mitochondrial membrane potential, reduces both mitochondrial and cellular ROS levels, and maintains cellular ATP homeostasis.

### NRP1 abrogation upregulates markers of cardiac stress and remodeling, while VEGF-B suppresses PGC1α Expression

To further analyze the effects of hypoxia on mitochondrial biogenesis and functions in H9c2 cells, we measured the mRNA expression levels of cardiac stress markers, such as *ANF* (*Atrial Natriuretic Peptide*), *Bnp* (*B-type Natriuretic Peptide*) and *βMHC* (*β myosine heavy chain)*, in H9c2 cells treated with hypoxia (CoCl_2_). In control normoxic cells, basal expression levels of *ANF, βMHC* and *BNP* were observed at moderate levels, reflecting normal mitochondrial function and low activation of cardiac stress markers **(Fig. 5A)**. However, upon hypoxia treatment, there was a significant increase in the expression of *ANF, βMHC,* and *BNP* **(Fig. 5A-5C),** indicating a response to hypoxic stress with the activation of mitochondrial biogenesis and increased cardiac stress signaling. The hypoxia-treated H9c2 cells suggested an adaptive response to mitochondrial dysfunction induced by low oxygen, while the increased expression of *ANF* and *BNP* may reflect myocardial stress.

**Figure 5.**
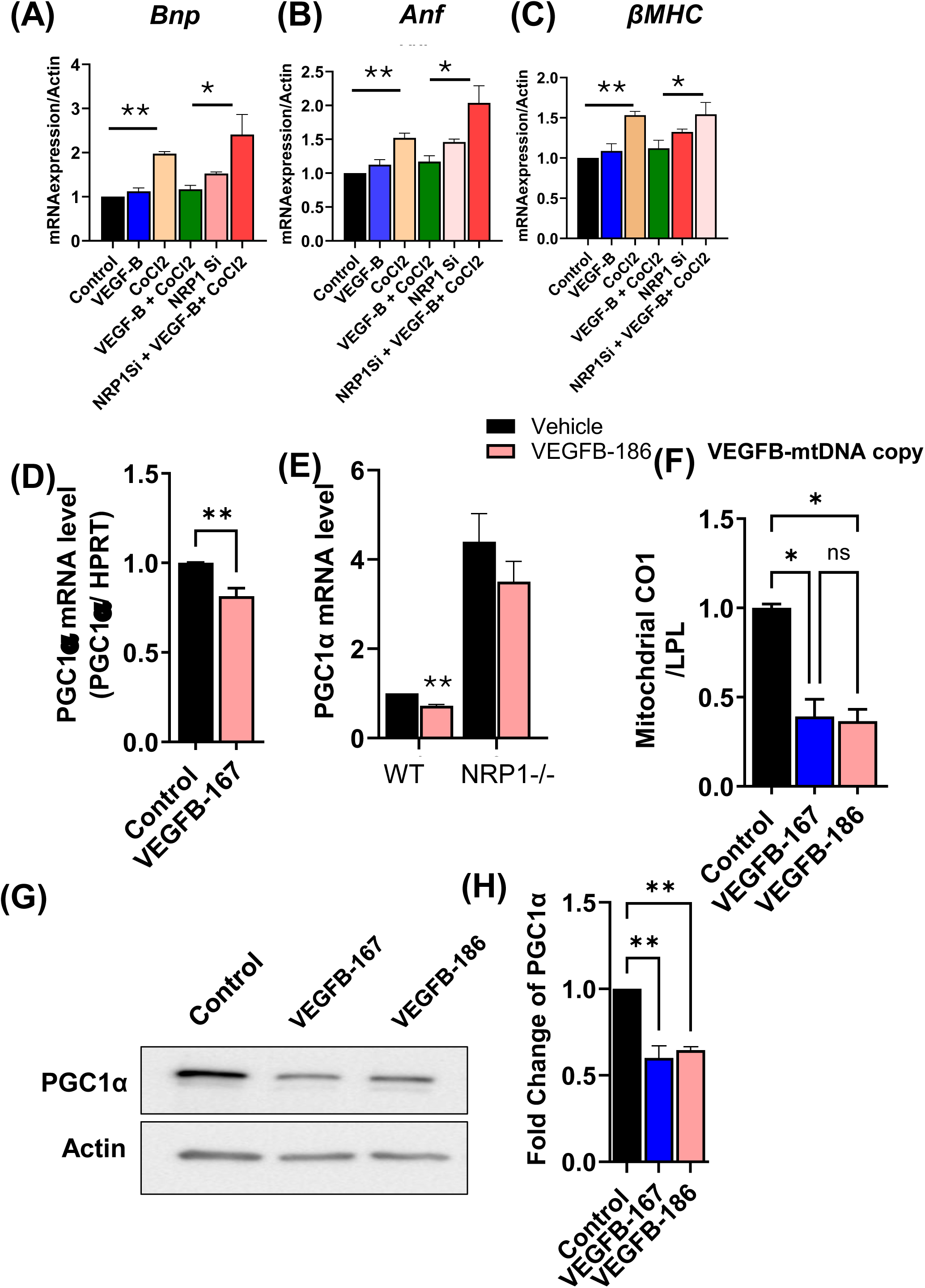
NRP1 Knockdown Blocks VEGFB-Mediated Regulation of Mitochondrial and Cardiac Remodeling Proteins. **(B)** Western blot analysis showing upregulation of the mitochondrial protein PGC1α and confirmation of NRP1 depletion in cardiomyocytes (CMs) isolated from neonatal NRP1 knockout (NRP1 KO) mice.(B) mRNA expression levels of cardiac remodeling genes in CMs from wild-type (WT) and NRP1 KO mice. **(C)** Relative mRNA expression of PGC1α in CMs isolated from WT and NRP1 KO mice following treatment with recombinant VEGFB. **(D)** Western blot analysis of PGC1α levels in CMs isolated from WT and NRP1 KO mice treated with GFP-AdV (control) or VEGFB-AdV (adenoviral overexpression), followed by VEGFB stimulation. **(E)** qRT-PCR analysis of ferroptosis markers in H9c2 cells. All RT-PCR experiments were performed in triplicate. Data are presented as mean ± SD. *p < 0.05,**p < 0.01.

Previous work by our group showed that NRP1 knockdown upregulates PGC1α expression, suggesting a role for NRP1 in repressing mitochondrial biogenesis pathways^35^. Interestingly, treatment with VEGF-B led to a marked reduction in PGC1α expression in wild-type CMs **(Fig. 5D)**, but this effect was abolished in the NRP1 knockdown CMs **(Fig. 5D)**, indicating an NRP1-dependent mechanism of VEGF-B action. Additionally, mitochondrial DNA (mtDNA) copy number was significantly decreased following VEGF-B treatment **(Fig. 5F)**, supporting the suppression of mitochondrial biogenesis. Wang et al showed that, NRP1 knockdown results in upregulation of cardiac remodeling genes such as ANF, BNP and βMHC, cMyc^35^. In our results, Consistent with this, we observed that the expression of ANF*, βMHC* and *BNP* remained comparable to that of hypoxia-treated cells, when compared to that NRP1 knockdown cells treated with VEGF-B and exposed to hypoxia, displayed confirmed that the loss of NRP1 disrupted the VEGF-B-mediated response.

### VEGF-B ameliorates the effect of hypoxia-induced stress in zebrafish hearts

The transparent nature of zebrafish embryos facilitates live imaging of cardiac function and vascular remodeling, making it an ideal system for studying VEGF-B-mediated cardio protection^36^. Notably, zebrafish are a valuable model for studying adult human heart disease due to their remarkable ability to regenerate cardiac tissue after injury, a capacity that is largely absent in adult mammals^37^. Despite structural differences, the zebrafish heart shares essential features with the human heart, including conserved cardiac genes, signaling pathways (e.g., Notch, Wnt, TGF-β), and electrophysiological properties^38^. Their genetic tractability, optical transparency (particularly in early life stages), and availability of transgenic reporter lines enable real-time imaging and precise manipulation of cardiac injury and repair processes^39^. These strengths make zebrafish an excellent system for modeling heart failure, cardiomyopathies, and ischemia, providing insights into mechanisms of cardiac remodeling and regeneration relevant to human cardiovascular disease. To further explore the protective role of VEGF-B/NRP1 signaling in cardiomyocyte survival and mitochondrial function under hypoxic stress, we developed a transgenic zebrafish model with inducible expression of VEGF-B (*Vegf-bb*) under the control of a heat-shock promoter driving the expression of VEGF-B: *Tg(*.pKTol2H70-LP2mc-zfvegfbb-gcG) adapted from a previous study by our lab^26^**(Fig. 6A)**. Using a cardiac-specific promoter, *Myosin Light Chain 7 (myl7)* or *Cardiac Myosin Light Chain 2 (cmlc2)*, we drive Cre expression exclusively in the zebrafish heart as described previously^40^. We generate a double transgenic *Tg(pKTol2H70-LP2mc-zfvegfbb-gcG);Tg(Cmlc2:CreER)* by crossing the *Tg(Cmlc2:CreER)* with the *Tg(*pKTol2H70-LP2mc-zfvegfbb-gcG fish. To confirm the expression of VEGF-B, 6–8-month-old adult zebrafish were exposed to a heat shock at 37°C for 30 minutes and 120 minutes, followed by tissue analysis. *VEGF-B* mRNA expression in the heart tissue isolated from control (-Cre) and the VEGF-B induced (+Cre) indicated strong *VEGF-B* expression in the heart tissues of the +Cre samples after both 30-minute and 120-minute heat shock exposures **(Fig. 6B)**. We observed a time-dependent increase in VEGF-B mRNA with high expression at 30 min, and reduced expression at 120-minute time. We then used ultra640-sound analysis to assess heart function by measuring ejection fraction (EF) and Fractional shortening (FS) from n=8 adult zebrafish as described before^41^ in the different groups following MI. The analyses were performed in a blinded manner where the group information was not available to the person performing the analysis. Both the control zebrafish and sham VEGF-B over-expressing zebrafish exhibited a normal ejection fraction **(Fig. 6C)** and fractional shortening **(Fig. 6D),** indicating healthy heart function. In contrast, MI in control (-Cre) fish showed a significantly reduced EF and FS compared to zebrafish without MI (sham) (50 % reduction) **(Fig. 6C**), consistent with impaired cardiac function due to myocardial injury. However, in the transgenic VEGF-B overexpressing zebrafish plus MI group, the ejection fraction was significantly improved after MI at 3 days post-MI compared to the control (-Cre) group, indicating a protective effect of VEGF-B on cardiac function. We then perform histological analysis using Hematoxylin and Eosin (H&E) staining and cleaved caspase-3 immunostaining to further analyze the beneficial effects of VEGF-B overexpression in zebrafish heart tissue sections. In the control group, H&E staining showed normal heart architecture with well-defined myocardial layers and no evidence of tissue damage **(Fig. 6E)**. In contrast, the MI group exhibited significant structural damage, including disorganized tissue structure, and extensive areas of necrosis and fibrosis **(Fig. 6E).** Cleaved caspase-3 staining in MI hearts also revealed immunoreactivity in the cardiomyocyte death, indicating widespread apoptosis **(Fig. 6F).** In the transgenic VEGF-B overexpressing zebrafish, H&E staining showed partial restoration of myocardial architecture, with a noticeable reduction in myocardial thinning and improved tissue organization after 3 days post MI with less evidence of fibrosis and necrosis compared to the control zebrafish MI group. Cleaved caspase-3 staining was also markedly reduced, and the number of apoptotic cells in the myocardial tissue was also reduced in the transgenic VEGF-B over-expressing zebrafish hearts after MI, compared to the control zebrafish MI group **(Fig. 6F)**, indicating that VEGF-B overexpression treatment effectively attenuated the hypoxia-induced cardiomyocyte apoptosis and promoted tissue preservation. These results collectively demonstrate that VEGF-B overexpression mitigates structural damage, reduces apoptosis, and improves cardiac function in a myocardial infarction model in transgenic zebrafish overexpressing VEGF-B, highlighting the therapeutic potential of VEGF-B in preserving heart tissue and function under MI stress responses. We next crossed the double transgenic fish with *Tg(hsp70;nrp1b;cmlc2-cas9-EGFP)*^41^ *to generate a triple transgenic line:Tg.HSP70:VEGF-B-eGFP);Tg(Cmlc2:CreER);Tg.HSP70:VEGF-B-eGFP);Tg(Cmlc2:CreER).* In this model, VEGF-B is overexpressed in the heart, and heat shock induces *nrp1b* knockout specifically in cardiomyocytes. Functional analysis of these triple transgenic fish revealed that *nrp1b* ablation abolishes the cardioprotective effects of VEGF-B, as heat-induced VEGF-B expression no longer improved cardiac function **(Fig. 6C, 6D)**. Histological analysis further supported this observation, showing a larger necrotic area post-injury in *nrp1b*-deficient hearts compared to VEGF-B overexpressing hearts with intact *nrp1b* **(Fig. 6E, 6F)**, indicating that NRP1 is essential for VEGF-B mediated protection.

**Figure 6.**
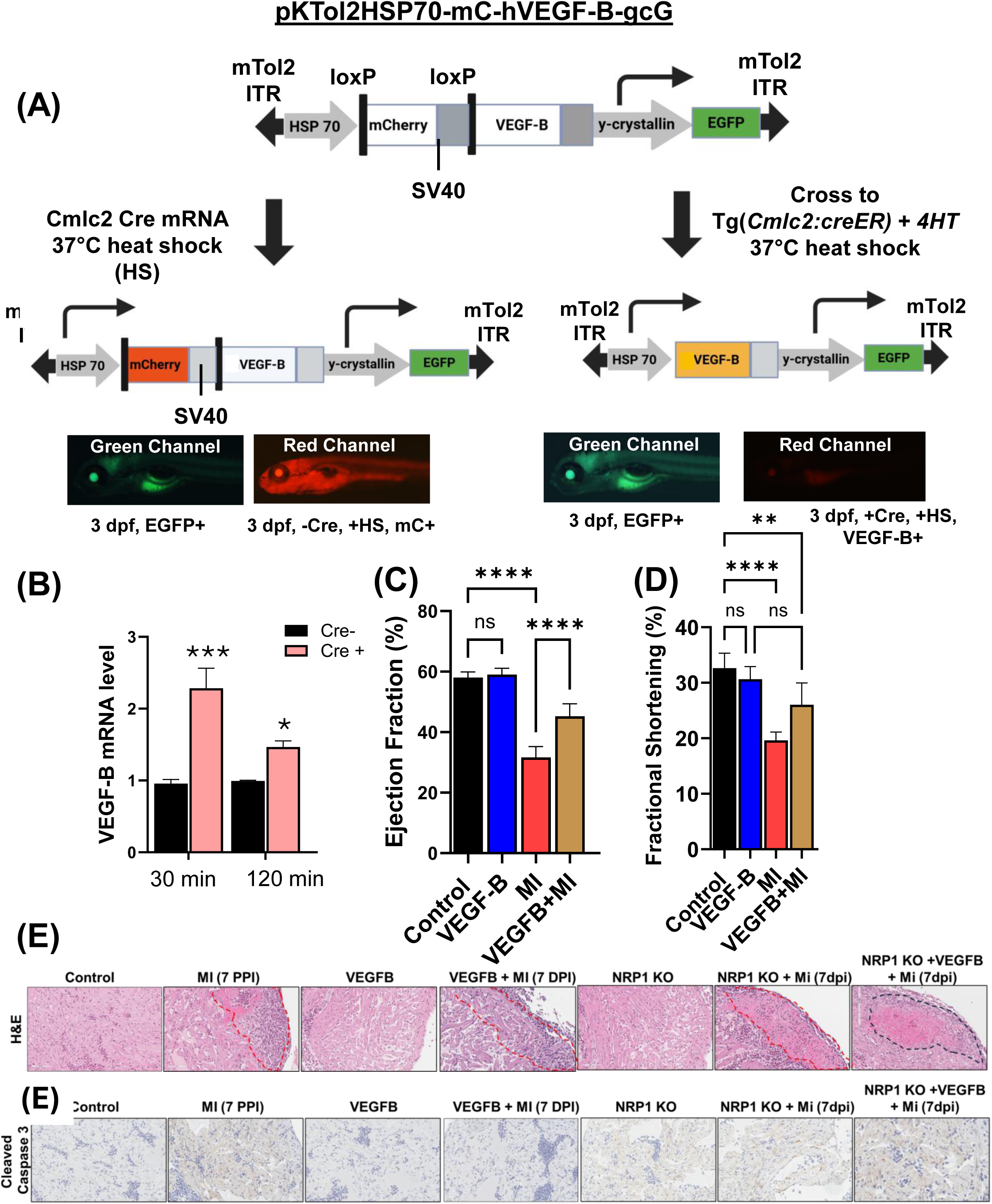
VEGFB mitigates the mitochondrial function and apoptosis in ischemic hearts in zebrafish. **(A)** The experimental outline shows the schematics of the generation of the heat shock inducible VEGF-B transgenic (The pKTol2H70-mC-hVEGF-B-gcG) composed of a heat-inducible HSP70 promoter driving a floxed mCherry gene and zebrafish VEGF-B (*vegf-bb*) and a γ-crystallin promoter driving EGFP. mTol2 ITR indicates mini-Tol2 plasmid inverted terminal repeat; BGH(A), bovine growth hormone polyadenylation signal; SV40(A), simian virus 40 polyadenylation signal; rβG(A), rabbit β-globin polyadenylation signal. The HSP70 promoter drives transcription of the mCherry gene, producing a red fluorescent protein in transgenic zebrafish. The lens-specific γ-crystallin promoter drives EGFP in the eyes. mC indicates mCherry; HS, heat shock. Crossing with Tg(cmlc2:creER) results in the excision of the floxed mCherry gene, and subsequent heat-shock induction of the HSP70 promoter produces hVEGF. Non-crossed transgenic zebrafish were heat shocked at 37°C at 3 dpf to monitor the temporal expression of mCherry and hVEGF. **(B)** mRNA expression of VEGFB across time after heat shock. **(C)** Heart function analysis of zebrafish after MI injury and VEGFB overexpression. **(D)** Histological analysis shows an H&E-stained heart section. **(E)** VEGFB-mediated enhancement of apoptotic heart as indicated by cleaved caspase-stained heart section after MI induction and VEGFB overexpression. The analysis was performed 7 days post MI induction. N=3 different sections from different animals were analyzed. * p<0.05, and **, p<0.01. ***, p<0.001.

### Spatial Distribution and Functional Insights of VEGF-B and NRP1 in Heart Cell Types

To examine the expression of VEGF-B and NRP1 in healthy and diseased heart tissue, we first used silico analysis to determine their differential expression patterns. UMAP plots generated from the Human Protein Atlas (HPA) database revealed that both VEGF-B and NRP1 exhibited prominent expression in heart muscle cells, with VEGF-B showing higher expression in cardiomyocytes and fibroblasts, while *NRP1* was most strongly expressed in endothelial cells **(Fig. 7A)**. The UMAP plots also indicate their tissue-specific expression patterns. The clusters c0, c1, and c5 corresponded to cardiomyocytes that exhibited higher VEGF-B expression than the other clusters. Similarly, for *NRP1*, the UMAP plot **(Fig. 7B)** showed the RNA single-cell expression clusters in the normal heart muscle, extracted directly from the HPA database. Clusters c3, c4, and c6 demonstrated high NRP1 expressions. Cluster c3 was identified as smooth muscle cells, cluster c4 as endothelial cells, and cluster c6 as another subset of endothelial cells. These results align with previous studies showing that *NRP1*, a cell surface receptor that binds to *VEGF*, plays a role in endothelial permeability, chemotaxis, and proliferation, as well as in the function of class III semaphorins^21,42,43^.

**Figure 7.**
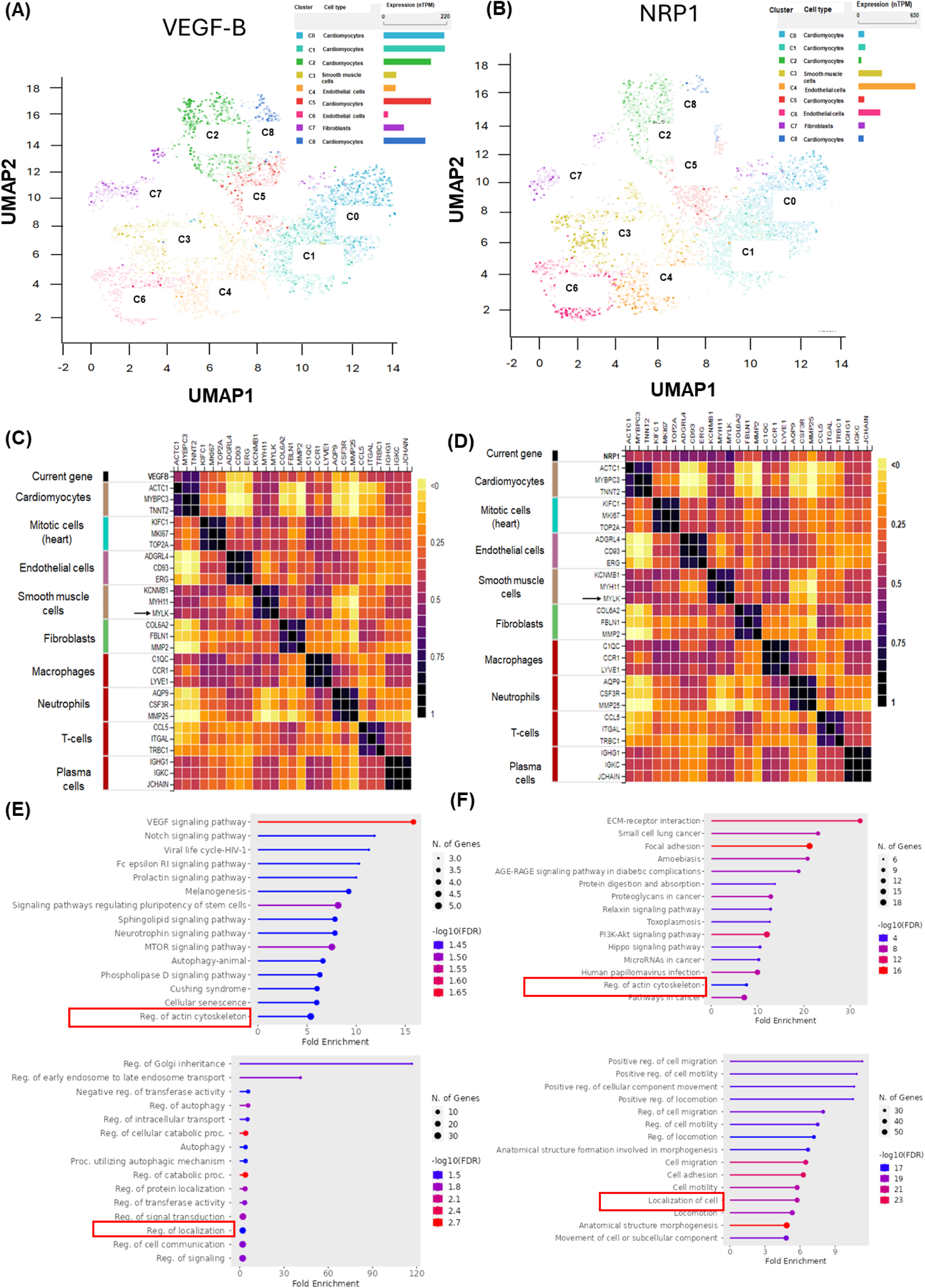
Single-Cell Expression Profiles and Pathway Enrichment of NRP1 and VEGF-B in Normal Heart Muscle. **(A)** UMAP plot showing the cell clustering in normal heart muscle tissue (Human Protein Atlas, HPA). Each dot represents a cell; clusters are color-coded with percentages based on the HPA method. **(B)** UMAP plot of NRP1 expression across cardiac cell clusters (HPA). **(C)** Heat map of tissue-specific expression of VEGFB in different cardiac cell types (HPA). Data are plotted as Z-scores normalized to mean 0 and standard deviation 1. **(D)** Heat map of tissue-specific expression of NRP1 in different cardiac cell types (HPA). **(E)** Lollipop plot displaying the top 50 enriched pathways from the 100 genes co-expressed with VEGFB. X-axis shows fold enrichment; circle size represents the number of genes per pathway; color indicates –log10(FDR). **(F)** Lollipop plot displaying the top 50 enriched pathways from the 100 genes co-expressed with NRP1, with the same visual parameters as in (E).

### Cell-Type Marker Analysis and Heatmap Visualization

To further explore the tissue-specific expression of VEGF-B and NRP1, we generated a heatmap visualizing their expression in conjunction with well-established cell-type markers in heart tissue. The heatmap revealed that VEGF-B was highly expressed in cardiomyocytes represented by markers such as *ACTC1* (*Actin alpha cardiac muscle 1*), *MYBPC3* (*Myosin binding protein C3*), and *TNNT2* (*Troponin T2*), fibroblasts, and smooth muscle cells, whereas *NRP1* showed strong expression in endothelial cells **(Fig. 7C, 7D)**. Specifically, genes such as *ADGRL4* (*adhesion G protein-coupled receptor L4*) from the endothelial cluster and *MYLK* from the smooth muscle cell cluster showed a strong positive correlation with NRP1 expression. **(Fig. 7D)**.

### Pathway Enrichment Analysis

We performed enrichment analysis using the ShinyGO 0.8 tool. The top10 enriched pathways are presented in (**Fig. 7E, 7F and Fig. S7)**. For VEGF-B, the most enriched pathways include VEGF signaling (Fold enrichment= 17, FDR <0.05), Notch signaling pathway (Fold enrichment= 12.8, FDR<0.05), Phospholipase signaling (Fold enrichment= 6.79, FDR<0.05), Regulation of actin cytoskeleton (Fold enrichment=5.79, FDR<0.05). Similarly, for NRP1 we observed Focal adhesion (Fold enrichment= 21.9, FDR<0.05), ECM-receptor interaction (Fold enrichment=33.19, FDR<0.05), Small cell lung cancer (Fold enrichment= 23.81, FDR<0.05) and Regulation of actin cytoskeleton (Fold enrichment= 7.85, FDR<0.05) as most enriched pathways besides Relaxin signaling, PI3K-AKT Signaling, AGE-RAGE signaling. Our analysis revealed that both VEGF-B and NRP1 might regulate pathways involved in Focal adhesion, actin cytoskeleton regulation, and cell and cell junction pathway localization **(Fig. 7E, 7F and Supplementary Fig. S7)**. The top 100 pathways for both VEGF-B and NRP1, along with the corresponding genes and enrichment scores, are presented in detail in (**Supplementary Tables S2 and S3).** The single cell RNA expression profiles of VEGF-B (median nTPM=217 in 429 sample) and NRP1 (median nTPM = 76.8 in 432 samples) across different cell types (tissues) in healthy humans is presented in (**Supplementary Fig. S8 and S9).**

## Discussion

VEGF-B plays a pivotal role in cardiovascular health, particularly in modulating mitochondrial function and cellular survival under stress conditions such as ischemia^16,44^. VEGF-B, through its receptor NRP1, interacts with the VEGF receptor 1 (VEGFR1), initiating signaling cascades essential for angiogenesis, tissue protection, and metabolic regulation^5,45,46^. NRP1, as a co-receptor, NRP1 facilitates VEGF-B signaling in endothelial cells and cardiomyocytes, enhancing its protective effects in the cardiovascular system^16^.

Our findings demonstrate that via the NRP1 signaling, VEGF-B enhances mitochondrial function and promotes cardiomyocyte survival under ischemic injury. Given the central role of mitochondria in energy metabolism and apoptosis, their preservation during ischemia is essential. Under both 1% O₂ and CoCl₂-induced hypoxic conditions, VEGF-B treatment mitigated cellular injury by reducing ROS, increasing ATP production, and decreasing apoptosis. These findings suggest that VEGF-B preserves mitochondrial function and supports cellular energy homeostasis under hypoxic stress. Moreover, VEGF-B’s impact on mitochondrial resilience under hypoxic stress points to its broader role in cardio protection. Despite VEGF-B’s limited angiogenic potency compared to VEGF-A, its impact on mitochondrial integrity and metabolic adaptation highlights its critical role in cardioprotection^47^.

Mechanistically, VEGF-B/NRP1 signaling appears to support mitochondrial efficiency, potentially through modulation of oxidative stress and enhancement of electron transport chain (ETC) activity. The AKT pathway is a key regulator of cellular survival, growth, and metabolism, and its activation by VEGF-B is associated with improved cellular resilience to stress, particularly in the context of cardiovascular injury^48^. Similarly, the ERK pathway, known for regulating cell proliferation and survival, suggests that VEGF-B may aid in vascular remodeling and endothelial cell protection^14^. Our results show that VEGF-B treatment activates both AKT and ERK signaling pathways, further supporting their pro-survival and anti-apoptotic roles in the CMs. These findings highlight VEGF-B’s dual role in both endothelial cell function and myocardial cell survival under ischemic conditions.

VEGF-B’s involvement in lipid metabolism and its regulation of mitochondrial dynamics may provide new insights into the therapeutic potential of VEGF-B in addressing metabolic dysfunctions that often accompany cardiovascular diseases^44^. By enhancing mitochondrial function, VEGF-B could promote better energy substrate utilization and optimize cellular responses to ischemic injury. In heart failure (HF), ferroptosis leads to substantial impairments in CMs, making targeted therapies against oxidative stress and ferroptosis an important option in HF research^49^. Nevertheless, the effective delivery of these therapeutic molecules and maximizing their efficacy remain significant challenges in current research efforts. Our data confirmed that VEGF-B effectively attenuated hypoxia-induced apoptosis and ferroptosis in cardiomyocytes. Beyond apoptosis, VEGF-B treatment also attenuated hypoxia-induced ferroptosis in cardiomyocytes, which is increasingly recognized in the pathogenesis of HF. As ferroptosis contributes to cardiomyocyte loss in HF, targeting this pathway via VEGF-B/NRP1 signaling presents a novel therapeutic avenue.

Furthermore, we observed upregulation of PGC-1α in NRP1-deficient cells, suggesting a compensatory response to mitochondrial dysfunction. Conversely, VEGF-B treatment reduced PGC-1α expression under normoxia, consistent with improved mitochondrial homeostasis^35^. Increased expression of stress markers ANF and BNP under hypoxic conditions aligned with myocardial stress responses and were attenuated by VEGF-B. In VEGF-B treated hypoxic cardiomyocytes, accompanied by enhanced pAKT and pERK signaling and decreased cleaved caspase-3 expression, indicating reduced apoptosis. and VEGF-B mitigated oxidative stress by promoting survival signaling pathways. However, in NRP1 knockdown cells, VEGF-B failed to lower ROS or inhibit apoptosis, underscoring the essential role of NRP1 in mediating VEGF-B’s protective effects. In NRP1 knockdown cells, ROS levels were significantly higher compared to those in VEGF-B-treated cells, even in the presence of VEGF-B and hypoxia. The lack of NRP1 expression diminished VEGF-B’s ability to reduce ROS production. In NRP1 knockdown plus hypoxia cells, ROS levels were comparable to those in hypoxia-only treated cells, indicating that NRP1 is critical for VEGF-B-mediated reduction of oxidative stress. The elevated ROS levels were associated with increased cleaved caspase-3 expression, confirming that oxidative stress was not effectively alleviated, and apoptosis was promoted due to the absence of NRP1. In NRP1 knockdown cells treated with VEGF-B and hypoxia, ROS levels remained high, similar to the hypoxia only group, confirming that NRP1 is essential for VEGF-B’s ability to reduce oxidative stress under hypoxic conditions. The persistently elevated ROS levels were accompanied by increased cleaved caspase-3 expression, further supporting the loss of VEGF-B’s protective effects due to NRP1 knockdown.

Importantly, to validate the protective role of VEGF-B in mitigating hypoxia-induced myocardial injury and preserving cardiac function, we developed a transgenic zebrafish model with inducible, cardiomyocyte-specific VEGF-B expression using the Cre-loxP system. Heat shock induction successfully activated VEGF-B expression, enabling controlled temporal studies^26^. In a zebrafish myocardial infarction (MI) model, VEGF-B significantly improved cardiac function, as evidenced by increased ejection fraction and fractional shortening on ultrasound. Histological analysis revealed reduced myocardial thinning, fibrosis, and necrosis, along with improved tissue organization and suppressed apoptosis. These results highlight the potential of VEGF-B as a therapeutic strategy for preserving cardiac structure and function during ischemic injury in vivo.

Finally, integration of transcriptomic data from GTEx, the Human Protein Atlas, and GEO confirmed enriched expression of VEGF-B and NRP1 in endothelial cells and cardiomyocytes, supporting their physiological role in maintaining cardiac homeostasis. In summary, our study identifies VEGF-B as a key regulator of mitochondrial function, redox balance, and survival signaling in cardiomyocytes under ischemic stress. By activating the AKT and ERK pathways and mitigating both apoptosis and ferroptosis, VEGF-B confers significant cardioprotection. These findings support the therapeutic potential of targeting the VEGF-B/NRP1 axis in myocardial infarction and heart failure. Further investigations are warranted to fully delineate the mechanistic underpinnings and to explore clinical applications.

## Conclusion

In this study, we elucidated the role of VEGF-B/NRP1 signaling in regulating mitochondrial function and ferroptosis in cardiomyocytes under hypoxic stress. Our findings demonstrate that VEGF-B enhances mitochondrial homeostasis by reducing ROS accumulation, increasing ATP production, and limiting apoptosis, thereby promoting cellular survival. These protective effects are mediated through NRP1, establishing VEGF-B/NRP1 as a critical axis in cardiomyocyte resilience to ischemic injury.

Furthermore, the ability of VEGF-B to mitigate ferroptosis underscores its broader role in modulating cell death pathways beyond apoptosis. Collectively, our results highlight VEGF-B as a promising therapeutic target for myocardial injury driven by mitochondrial dysfunction, warranting further investigation in preclinical and clinical models of cardiac disease.

## Acknowledgments

The authors thank the members of the Mukhopadhyay lab for their valuable input and suggestions. R.S.A acknowledges the support from the Innovation in Aging grant. The authors thank Dr. Karl J Clerk, Mayo Clinic, Rochester for providing the vector pKTol2 backbone.

## Author’s Contribution

R.S.A, and S.M.V wrote the original manuscript; R.S.A, S.V.M, R.K, C.M, A.D, A.S, E.W and S.D performed the in vitro and in vivo studies. Y.W helped with in vitro experiment. D.M and R.S.A conceptualized the study. S.E and A.S helped in generating the VEGF-B construct. D.M acquired funding, and D.M and Y.W supervised the work.

